# Differential Proteomic Analysis of DEN-Induced Hepatocellular Carcinoma in Male and Female Balb/c Mice Reveals Novel Gender Specific Markers

**DOI:** 10.1101/2025.05.15.654000

**Authors:** Salmma Salamah Salihah, Muhammad Tahir, Bareera Bibi, Rabia Sultan, Martin R. Larsen, Munazza Raza Mirza, Sana Mahmood, Muhammad Rizwan Alam, Jamila Iqbal, Asma Gul

## Abstract

**Background:** Hepatocellular carcinoma (HCC) is one of the leading causes of hepatic malignancy with a higher prevalence in males compared to females, however, the distinct underlying mechanisms contributing to this disparity remain poorly understood.

**Methods:** In this study, we aimed to investigate comparative proteome profiling of a diethylnitrosamine (DEN) induced HCC model in male and female Balb/c mice. We extracted proteins from liver tissue of DEN treated male and female mice and their corresponding controls and subjected them to mass spectrometry and subsequent bioinformatics analyses.

**Results:** We identified 170 and 233 differentially expressed proteins (DEPs) in female and male mice, respectively. We identified chemical carcinogenesis and cortical cytoskeleton as the shared pathways between the two groups. In addition, we identified distinct signaling pathways in DEN-treated male and female mice. Female mice showed enrichment in oxidative phosphorylation, fatty acid biosynthesis, metabolism and degradation and cytochrome P450 clusters. In contrast, in male mice, these pathways were enriched in cholesterol metabolism, coagulation and mRNA processing. Further, we identified top ten genes ranked by highest maximal clique centrality, by protein-protein interaction analysis of differentially expressed proteins (DEPs) in both sexes. Of these hub genes, female mice showed upregulation of NDUFA8 and ATP5H and were associated with poor patient survival. On the other hand, In DEN-treated male mice upregulation of FGG, FGA, HPX and SERPINC1 were associated with poor survival.

**Conclusion:** In conclusion, our research provides gender-specific proteomic signatures in DEN-induced HCC. The identification of proteins associated cholesterol metabolism and coagulation in males, and mitochondrial complex I proteins in females as prognostic markers suggests novel therapeutic targets that may inform gender-tailored treatment strategies for HCC.

**Simple Summary:** Hepatocellular carcinoma (HCC) is a common and deadly liver cancer that affects men more than women. To understand the biological reasons behind this difference, we developed a liver cancer model in male and female mice using a chemical called diethylnitrosamine (DEN). We then studied the proteins involved in tumor development using advanced techniques like mass spectrometry and bioinformatics. We found that different sets of proteins and biological pathways were active in males and females. In males, ribosomal and RNA-binding proteins were linked to worse survival, while in females, mitochondrial proteins were more important. These findings suggest that men and women may need different strategies for diagnosing and treating HCC, and they offer new gender-specific targets for future therapies.

## 1. Introduction

Hepatocellular carcinoma (HCC) remains one of the most prevalent malignancies worldwide [1], posing a great burden on global health [2]. Notably, significant sex-based differences exist in susceptibility and survival outcomes: males are more prone to hepatocarcinogenesis [3], while females often demonstrate better prognoses [4]. These differences suggest underlying biological and molecular mechanisms that remain poorly understood, especially at the proteomic level. Investigating these mechanisms could unlock new opportunities for precision medicine and sex-specific treatment approaches.

Animal models are central to preclinical research in HCC, offering crucial insights into the molecular and pathophysiological landscape of the disease. While gene-editing and in-situ transplantation models allow focused mechanistic studies, they often fall short in capturing the systemic tumor-immune dynamics seen in human disease [5]. In contrast, the diethylnitrosamine (DEN)-induced mouse model closely mirrors the transcriptomic and proteomic characteristics of aggressive human HCC, particularly the poor-prognosis S-III subtype [6]. It replicates key features such as low β-catenin expression, high proliferation, chromosomal instability, and altered apoptotic signaling [7]. This makes the DEN model highly translational and suitable for investigating complex interactions in hepatocarcinogenesis, including potential sex-based differences. Two relevant studies have advanced this space. Zhang et al. [8] profiled the DEN and CCl_4_-induced male mouse model and aligned it with human HCC subtypes, identifying prognostic biomarkers. Rong et al. [9] explored gender-based proteomic differences in Ras-oncogene–induced HCC, finding that male hepatocytes were more vulnerable to oncogenic stress.

Despite these advances, critical gaps remain in understanding sex-specific molecular mechanisms in HCC. Zhang et al.’s study focused exclusively on male mice, leaving the female proteomic landscape unexamined [8]. While Rong et al. shed light on sex-related vulnerability using a Ras-driven model, their findings were model-specific and may not generalize to chemically induced HCC [9]. As a result, there is a lack of comprehensive, comparative proteomic data on male and female responses in a highly translational model like DEN.

In this study, we conducted a comprehensive proteomic analysis using a diethylnitrosamine (DEN)-induced mouse model of hepatocellular carcinoma (HCC), directly comparing tumor profiles between male and female subjects. This model, known for its strong translational relevance to aggressive forms of human HCC, enabled us to investigate sex-specific molecular differences in tumor biology. Our analysis revealed several key findings: (1) distinct proteomic signatures unique to male and female HCC, (2) pathways specifically involved in male and female hepatocarcinogenesis, (3) male and female-specific proteins associated with poorer outcomes, and (4) candidate therapeutic targets for developing sex-tailored treatment strategies. These results enhance our understanding of the molecular underpinnings of HCC and provide a foundation for future precision oncology approaches that consider biological sex as a critical variable.

## 2. Materials and Methods

### 2.1. Mouse model and tissue collection

Balb/c mice were obtained from the National Institute of Health, Islamabad, Pakistan. All procedures were approved by the Ethical Committee, of the International Islamic University Islamabad, and carried out in accordance with institutional animal care guidelines. A total of 21 mice (11 females and 10 males) were housed under standard laboratory conditions with a 12-hour light/dark cycle and a controlled temperature of 22–25°C. Animals had free access to a standard diet and water throughout the study. The mice were randomly divided into control and treatment groups, with each group consisting of 5 males and 6 females. At four weeks of age, mice in the treatment group received a single intraperitoneal injection of diethylnitrosamine (DEN) (100 mg/kg body weight, Sigma-Aldrich, Cat # N0756), while control animals were injected with an equivalent volume of normal saline. Both groups were monitored for 32 weeks.

At the end of the experimental period, surviving animals were terminally anesthetized with an overdose of sodium pentobarbital (Sigma-Aldrich, Cat # P3761). Liver tissues were collected from all animals and rinsed in phosphate-buffered saline (PBS) (Sigma-Aldrich, Cat # P4739). Each liver was examined macroscopically for visible nodules or lesions, then sectioned for further analysis. One portion was fixed in 4% formaldehyde (Sigma-Aldrich, Cat #252549) for histological and immunohistochemical evaluation, while the remaining tissue was stored for subsequent proteomic analyses.

### 2.2. H&E staining & immunohistostaining

Formaldehyde-fixed tissues were embedded in paraffin and sectioned into 5um slices. These sections were mounted onto glass slides and deparaffinized using three changes of xylene. Rehydration was performed through a graded ethanol series (100%, 95%, and 70%), after which the slides were stained with Mayer’s hematoxylin for 5 minutes and counterstained with eosin. Sections were then dehydrated through an ascending ethanol series (70%, 95%, and 100%), followed by clearing in xylene. The stained slides were mounted using Permount and allowed to dry before microscopic evaluation.

For immunohistochemical analysis, antigen retrieval was performed using a microwave-based method in 0.01 M sodium citrate buffer (pH 6.0), as previously described by Maffini et al. [10]. The following primary antibodies were used: Vimentin (Cat # NCL-VIM-V9, Leica Biosystems; 1:1000), Ki 67 (Cat # PA0118, Leica biosystems) and E-Cadherin (Cat # PA0387, Leica Biosystems; 1:1000–1:4000). Antigen–antibody complexes were visualized using a streptavidin–peroxidase detection system with diaminobenzidine (DAB; Sigma-Aldrich) as the chromogen. Slides were counterstained with Harris’ hematoxylin. Microscopic images were captured using an Olympus BX53 microscope equipped with a digital Olympus camera.

### 2.3. Sample preparation for proteome analysis

Liver sections of Balb/c reserved for proteome analysis were rinsed with phosphate buffer saline (PBS) and immersed in freshly supplemented RIPA buffer (10 mM Tris-HCl, 140 mM NaCl, 0.1% SDS, 1% Triton X 100, and 0.01% Sodium deoxycholate) (R0278, Sigma Aldrich). The tissues were homogenized on ice for 20 minutes using a tissue homogenizer. After centrifugation (13, 000× *g*; 20 min, 4 °C), supernatants were stored at-80°C. Protein estimation was performed through Bradford Assay.

### 2.4. Tryptic digestion

For the enzymatic digestion of protein standard (a-casein, β-casein, and ovalbumin) and protein samples, 0.1mg/mL(200μL) was aliquoted. To adjust the pH (∼8) 1M NH_4_HCO_3_ (160μL) was added to the aliquots. Protein samples were reduced and denatured by adding 45 mM Dithiothreitol (DTT) and 40 mM nOGP. Afterward, the vials were incubated on a thermomixer (Eppendorf, Germany) for 30 mins at 90°C and 800 rpm. Protein solutions were allowed to cool at room temperature followed by alkylation at room temperature, carried out by adding 100 mM of iodoacetamide (IAA) and the vials were incubated in the dark for 15 mins. Deionized water was added followed by the addition of 2μg trypsin. After adding trypsin samples were digested on a thermomixer (Eppendorf, Germany) for 16 hours at 600 rpm and 37°C. The digestion process was stopped by adding 2% trifluoroacetic acid TFA (60μL, pH ≤3). All digests were stored at -20°C.

### 2.5. Mass spectrometry analysis

Peptide digests were analyzed by nanoLC-MS/MS system on a Q Exactive HF Orbitrap LC-MS/MS system (Thermo Fischer) coupled to an EASY-LC 1000 system (Thermo Fischer Scientific). Long-fused silica capillary column (PicoFirt 18 cm, 75 m inner diameter) was packed in-house with reversed-phase Repro-SilPur C18-AQ 3 m resin and used to load 1mg of purified peptides. Peptides were eluted from Buffer A (0.1% Formic acid) at a flow rate of 250 nL/min followed by elusion through buffer B (95% acetonitrile in 0.1% Formic acid). The gradient gradually increased in B from 5% to 28% in 40 mins, 45% in 60 mins, and finally to 100% in the end. The Q-Exactive HF was set up to operate in data-dependent acquisition mode with the following parameters: full scan automatic gain control (AGC) target 3e6 at 120000 FWHM resolution, scan range 350-1600 mz, Or-bitrap full scan maximum injection time 100 ms, normalized collision energy 28, a dynamic exclusion time of the 30s, an isolation window of 1.2 m/z and top 20 MS2 scans per MS full scan.

### 2.6. Secondary data acquisition

To compensate for the early loss of male mice, male cohort was supplemented with secondary data containing four DEN + CCl_4_ treated and four control male mice from a closely related study by Zhang et al. [8] to make the total count 12 in the male cohort. The raw LFQ intensities were downloaded from iprox (Accession number: PXD049148) (https://www.iprox.cn/page/BWV016.html).

### 2.7. Statistical data analysis

The statistical analysis began by importing protein groups from MaxQuant. Initial pre-processing was performed in Perseus v.1.6.10.50 for both male and female cohorts, where potential contaminants, reverse database hits, and proteins identified only by site were removed, followed by log2 transformation.

For the female cohort, the filtered data was further filtered based on valid values, followed by median imputation of missing values. Imputed data was then subjected to significance testing.

For the male cohort, the primary and secondary datasets were merged on common proteins, followed by filtration based on valid values in each group and batch. Missing values were imputed using the median imputation, and batch effects were corrected using the ComBat empirical Bayes method implemented in R version 4.5.1(package sva). From this point onward, the ComBat-corrected, merged male dataset was used for subsequent analyses.

Principal component analysis (PCA) was performed to confirm clear separation of control and treated samples in male and female datasets. Significance testing of protein LFQ intensities was performed separately for male and female cohorts using two-sample t-test in Perseus v.1.6.10.50. Proteins with a p-value ≤ 0.05 and Abs (log FC)>=1 were considered significant. PCA plots, heatmaps and volcano plots were generated in R version 4.5.1using ggplot2.

### 2.8. PPI and hub proteins

Protein-protein interaction (PPI) network of differentially expressed proteins in each group was constructed using STRING DB (version 12.0) (https://string-db.org/). The PPI networks were visualized in Cytoscape (version 3.10.3) [11]. Functional modules were identified through MCODE plugin and top 3 modules were selected based on MCODE score in the Cytoscape [12].The analysis of the female group was carried out at default parameters (k-score = 2, cut-off degree = 2, max depth size = 100 and node score cutoff = 0.2) and analysis of the male group was also carried out at default parameters with slight modification (k-score = 2, cut-off degree = 2, max depth size = 100 and node score cutoff = 0.1) Maximal Clique Centrality (MCC) algorithm was used to select hub genes from the PPI network using the CytoHubba plugin of Cytoscape [13].

### 2.9. Gene enrichment analysis

The biological significance of each module was determined through Gene Ontology (GO), Kyoto Encyclopedia of Genes and Genomes (KEGG: https://www.kegg.jp/kegg/pathway.html), and Reactome (https://reactome.org/Pathway.html) enrichment analyses to determine specific processes, molecular functions, or cellular components represented within each module. Overrepresentation analysis of GO, KEGG, and Reactome terms was performed using the R package clusterProfiler, considering only terms with an adjusted p-value ≤ 0.05 as statistically significant. For visualization, redundant GO terms were removed and only top five enriched terms in each category were plotted for each module.

### 2.10. Survival analysis

The clinical value of differentially expressed proteins was estimated by converting 20 hub proteins to their human homologs using the “biomaRt” R package. Protein expression of human homologous proteins corresponding to DEPs was matched with overall survival using the UALCAN tool.

## 3. Results

### 3.1. DEN-Induced Mouse Model for Male and Female HCC

DEN-induced HCC mouse model for male and female Balb/c mice was established following one of the methods described by [14] with some modifications. All six female mice in the treatment group survived until 8 months whereas only 2 male mice survived in each group till the end. Subsequent analysis was performed with the surviving individuals of the groups. Normal and tumor tissues with internal and external controls were collected from control and treated mice of both genders separately for the histological and proteomic analysis (**Figure 1a**).

**Figure 1.**
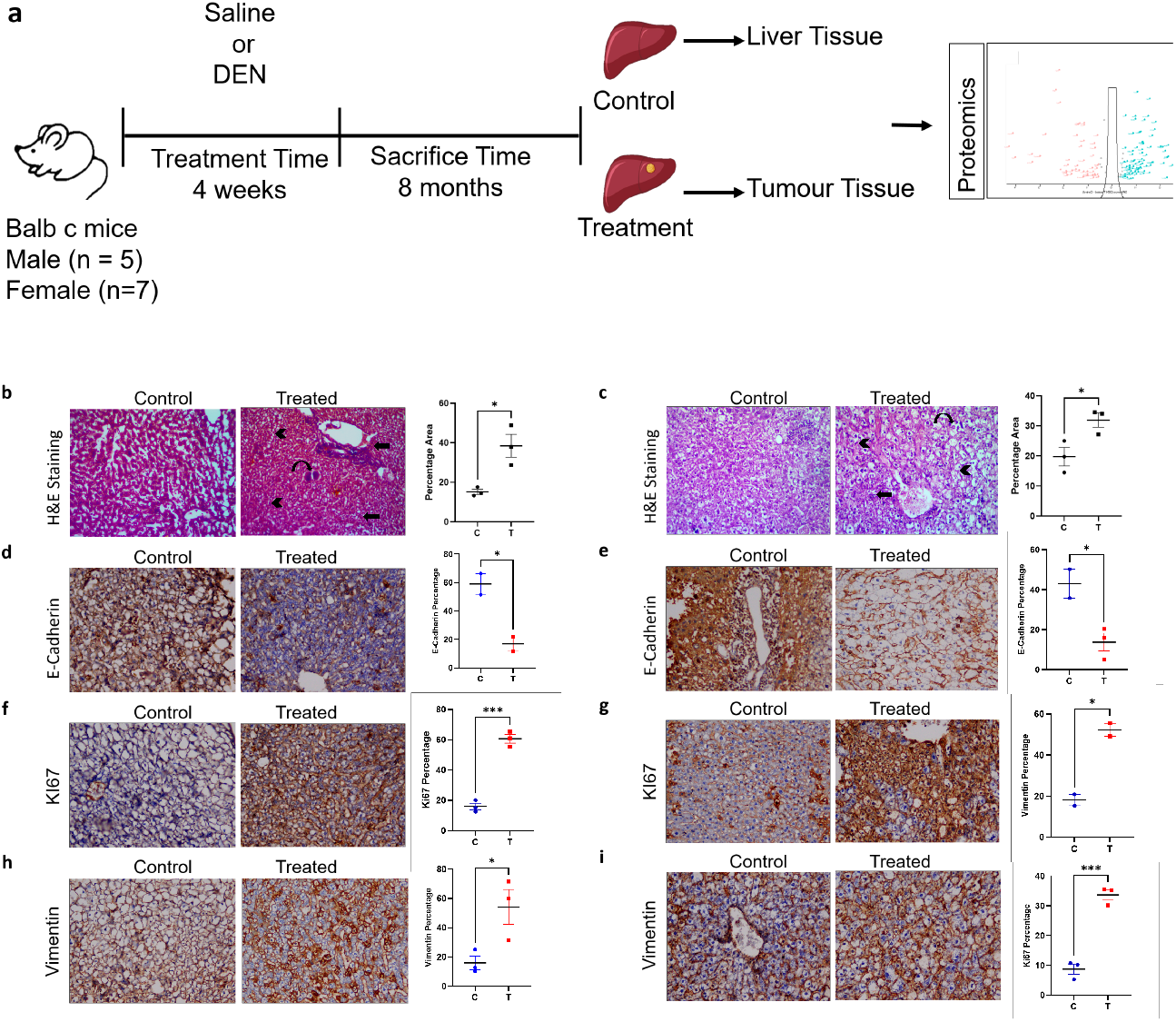
Schematic diagram of the study, H&E analysis and immunohistochemical staining of tissue sections from treatment and control groups (brown color indicates antigen specific immunostaining whereas blue color indicates nuclear hematoxylin staining); (**a)** Schematic diagram of the development of DEN-induced HCC in Balb/c mice; **(b, c)** H&E Staining (magnification 20X, 40X) of liver sections obtained from treated and control groups of male mice and female mice respectively. Half circular arrows in the treatment group indicate enlarged nuclei; arrows indicate immune infiltration and arrow heads show disorganized morphology. Percentage increase in the nuclear area is represented on the graph (n=3, n=3); (**d, e)** E-Cadherin staining showing loss of E-Cadherin in the treatment groups of male and female mice (n=2, n=3); (**f, g)** KI67 staining showing gain of KI67 in treated samples of male and female liver sections (n=3, n=3); (**h, i)** Vimentin staining showing increase in the vimentin expression in treated and control section of male and female mice liver (n=3, n=2).

In the DEN-induced mouse model multiple tumors developed in the liver. We compared the morphology of control and tumor tissue using H&E staining. Control mice showed neatly arranged hepatocytes whereas tumor tissue showed disorganized morphology, large nuclei, and immune infiltration suggesting DEN-induced carcinogenesis in the mouse liver. Percentage area of the nuclei increased from 15% to 38% and from 19 to 31% in male and female groups respectively (**Figure 1b, c**).

We also performed immunohistochemical analysis of tissue samples using tumor specific markers, E-Cadherin, vimentin and Ki67. The positive area stained with E-Cadherin decreased from 58 to 16% and 43 to 13% in treated samples of male and female mice respectively, suggesting loss of cellular adhesion (**Figure 2d, e**). Percentage area for the proliferation marker Ki 67 was increased from 15 to 60% and 8 to 33% in tumor samples of male and female samples respectively (**Figure 2f, g**). Increase in expression of Ki 67 shows increased cellular proliferation in treated samples. The positive area stained with vimentin was significantly increased from 16 to 54% and 28 to 48% in tumor tissue compared to normal tissues in male and female mice, respectively **(Figure 2h, i)**. These results indicated that the DEN-induced mouse model was successfully generated with characteristic proliferative and adhesive features of tumor.

**Figure 2.**
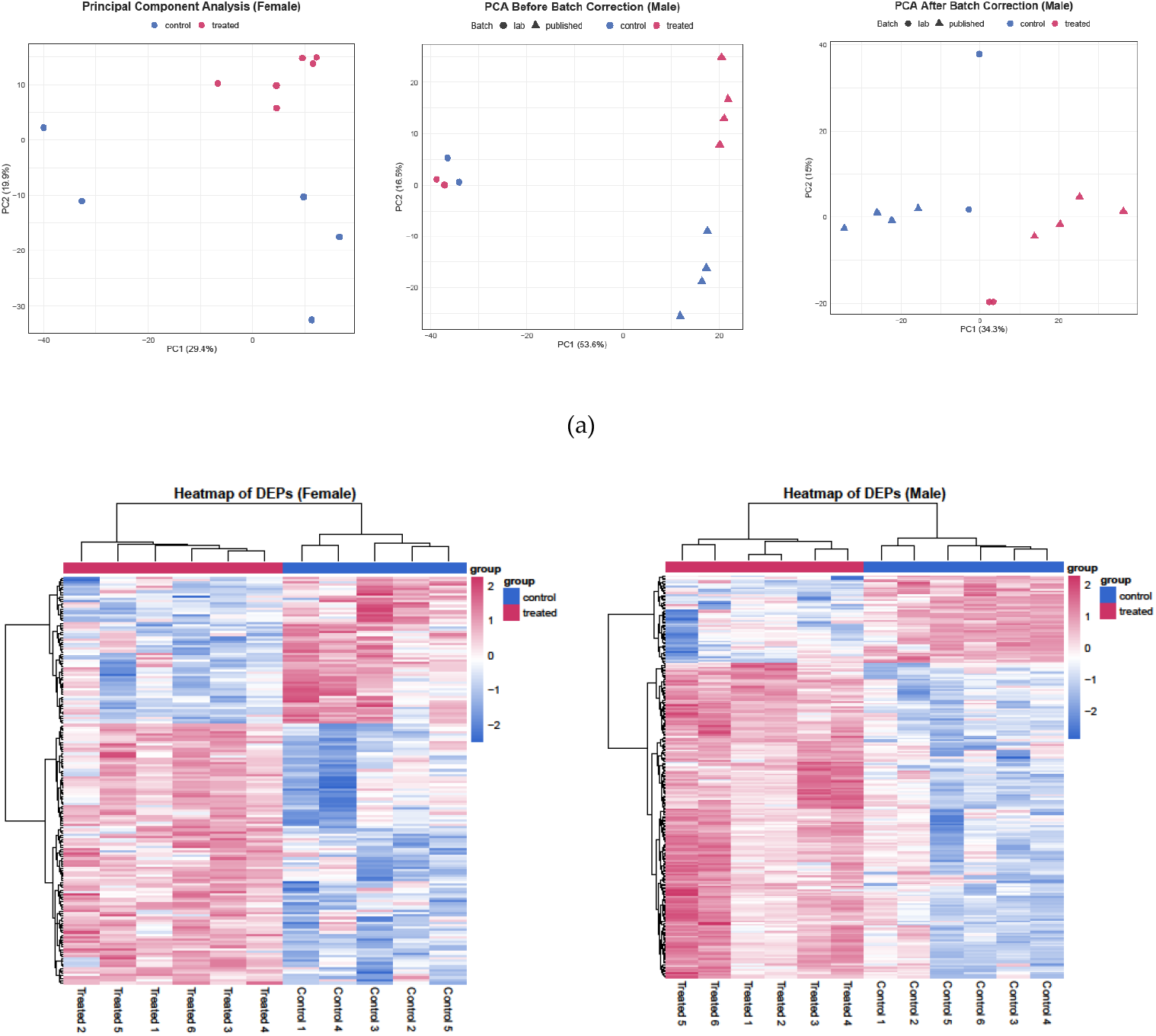
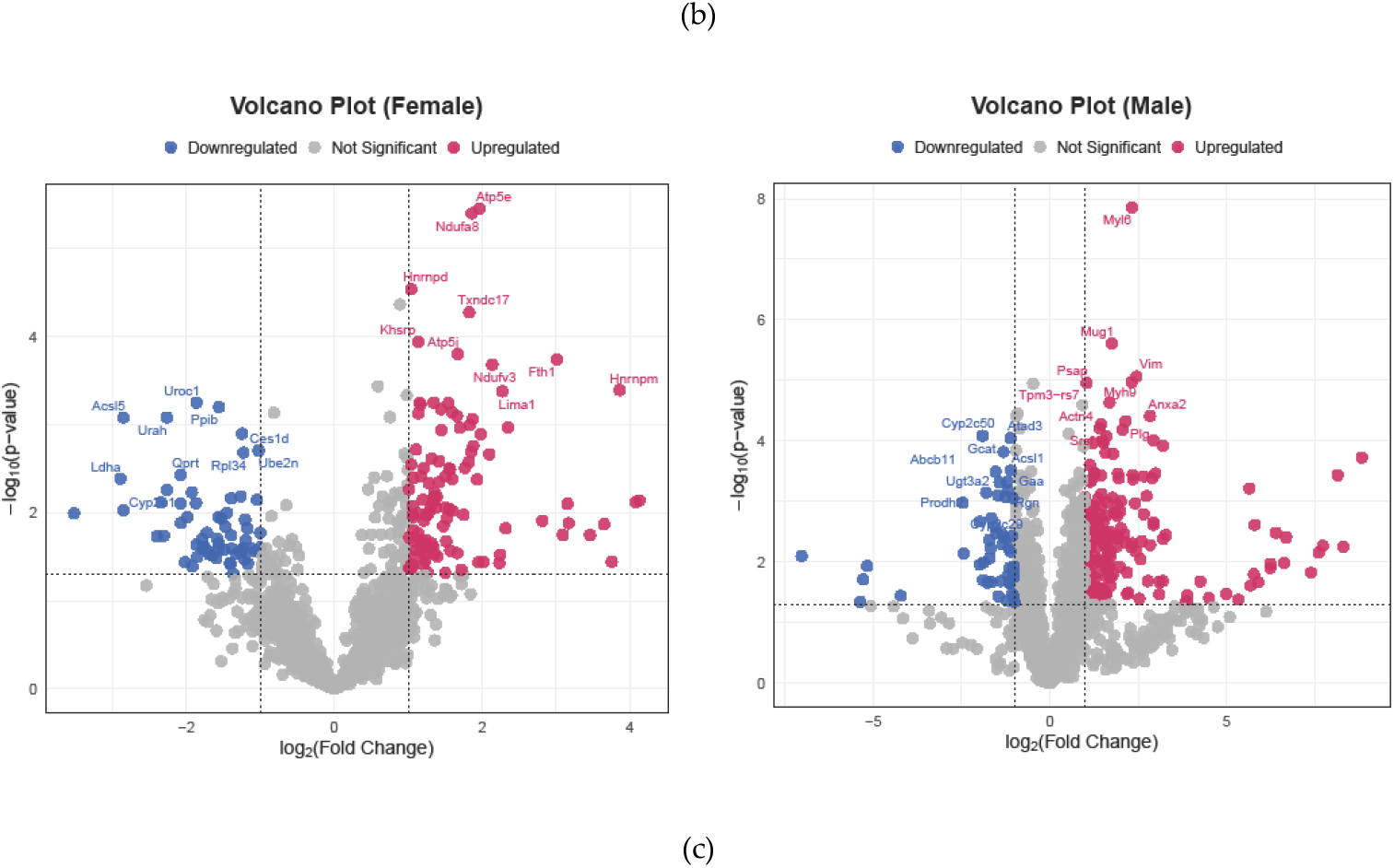
Proteomic analysis of DEN-induced HCC mouse models: **(a)** PCA represents clear separation between the treatment (red) and control (blue) groups of female mice, PCA of male cohort before and after ComBat indicate post ComBat separation of PC1 and PC2; (**b)**Heatmap of DEPs expression in liver tissue of treatment (red) and control groups (control) in both genders; (**c)** Volcano plots male of differentially expressed proteins in male and female mice. Blue dots represent downregulated proteins, red dots represent upregulated proteins, and grey dots represent proteins with insignificant differences (p-value < 0.05, |log2FC|> 1). All plots were created in R software version 4.5.1 (ggplot2).

### 3.2. Statistical data analysis

In the present study, protein samples isolated from 11 female and 4 male mice were analyzed through label-free LC-MS which led to the identification of total 2897 proteins in each sample. After the filtration of contaminants, only identified by site and reverse hits 2803 proteins were selected. For the female cohort, log 2 transformation followed by filtration in each group, reducing the number of proteins to 1232. Principal component analysis (PCA) revealed the spread of the data. Control and treated groups were clearly separated on the PCA plot for female mice with 29.4% contribution from component 1 and 19.9% contribution from component 2 (Figure 2a).

For the male cohort, after the filtration of potential contaminants, only identified by site and reverse database hits 2803 proteins were selected from the primary data whereas 3960 proteins were selected from the secondary data. The primary and secondary data sets were merged based on 2281 common proteins making the total number of samples 12 (6 treated and 6 control) in the male cohort. Log2 transformation followed by filtration in each group and batch, reduced the total number of proteins to 1377. Median imputation of missing values was carried out and the batch effects were corrected using ComBat batch correction (with group as covariate). Principal component analysis (PCA) after combat showed complete segregation of treatment and control groups (Figure 2a).

Significance of the expressed protein based on the difference between the LFQ intensities of treatment and control groups in both male and female cohort was calculated using two sample t test. In female mice, 170 proteins were significantly differentially expressed among which 62 proteins were downregulated (supplementary table 1) and 108 proteins were upregulated (supplementary table 2). In male mice, 233 proteins were found to be significantly differentially expressed among which 182 proteins were upregulated (supplementary table 3) whereas 51 proteins were downregulated (supplementary table 4). Heatmaps of the DEPs were plotted based on Euclidean distance in both male and female groups (Figure 2b). In addition, volcano plots for both groups with a cutoff (p-value < 0.05, |log2FC|>1) are shown in Figure 2c.

### 3.3. PPI networks in Male and Female mice

For DEPs in male and female mice, the PPI network was constructed using Cytoscape version 3.10.3. The female PPI network consisted of 143 nodes with 605 edges while the male PPI network consisted of 214 nodes with 1420 edges. MCODE score of the 1st module in the female group was 14 (14 nodes and 91 edges), whereas the score of second and the third group was 8.75 (9 nodes and 35 edges) and 8 (8 nodes and 28 edges) respectively (Figure 3a). MCODE score of first, second and third module in male group was 17.76 (18 nodes, 151 edges), 12.66 (13 nodes, 76 edges) and 9.83 (13 nodes, 59 edges) respectively. Ten highest ranked genes based on MCC algorithm were selected as hub genes45gg (Figure 3b).

**Figure 3.**
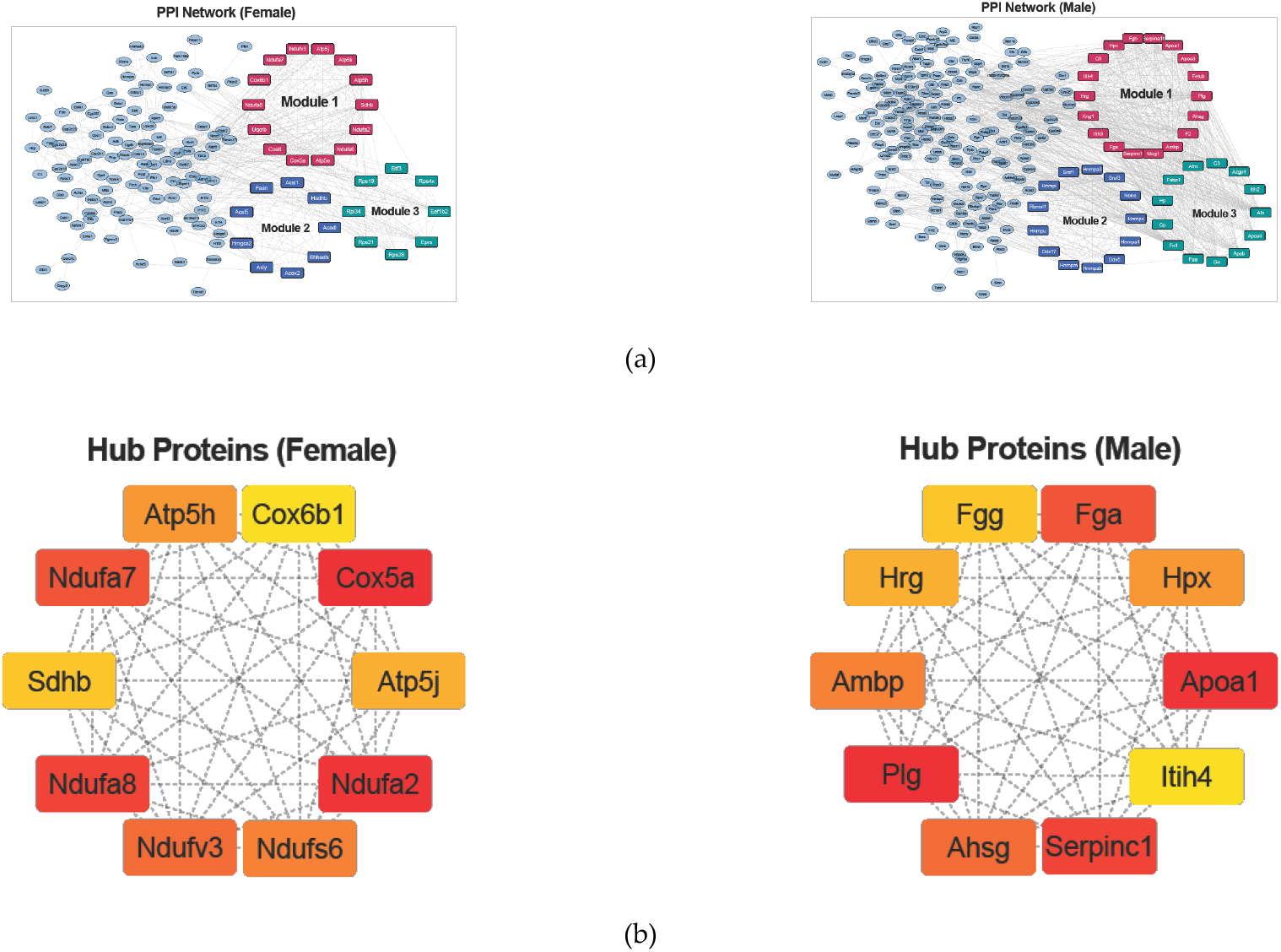
Protein-protein interaction in the differentially expressed proteins of both genders: **(a)** Pathway interaction in the top three modules of male and female DEPs; (**b)** Hub protein analysis of the top cluster.

### 3.3. Oxidative phosphorylation and fatty acid metabolism in Females

Functional enrichment analysis Gene Ontology (GO), Reactome Analysis, and KEGG pathway analysis) of top 3 modules was performed to study the biological relevance of significantly altered proteins. GO analysis of selected modules revealed significant enrichment in metabolic processes associated to thioester, fatty acid, acyl-CoA, ATP, ribose phosphate, rRNA and cellular lipid catabolism. Processes related to the biogenesis, assembly and maturation of SSU™rRNA and structural constituent of ribosome rRNA binding were also enriched. Moreover, DEPs related with mitochondrial structure and respiration such as respiratory chain complex, mitochondrial respirasome, mitochondrial protein™containing complex, electron transfer activity and proton motive force™driven mitochondrial ATP synthesis were also enriched (Figure 4a).

**Figure 4.**
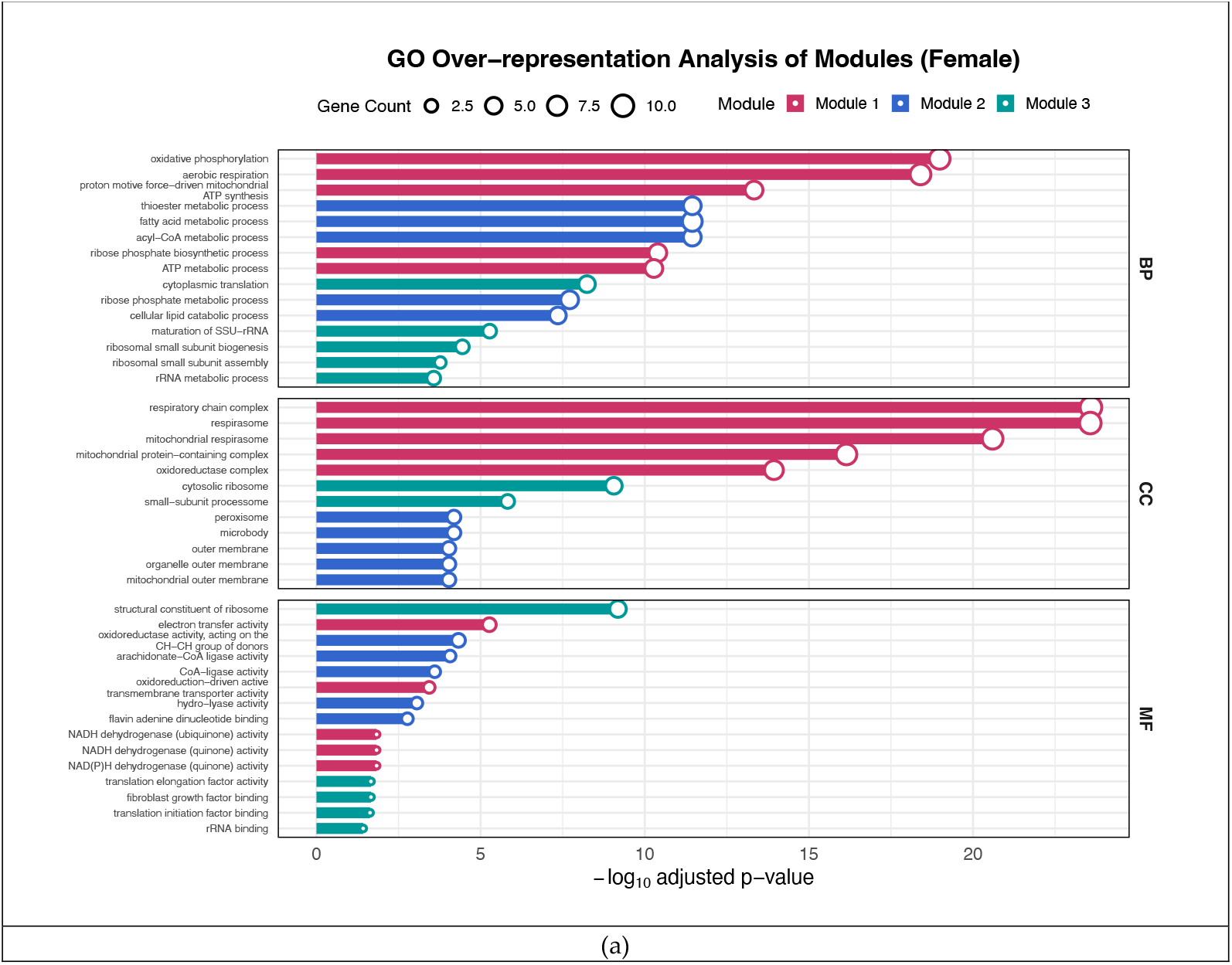
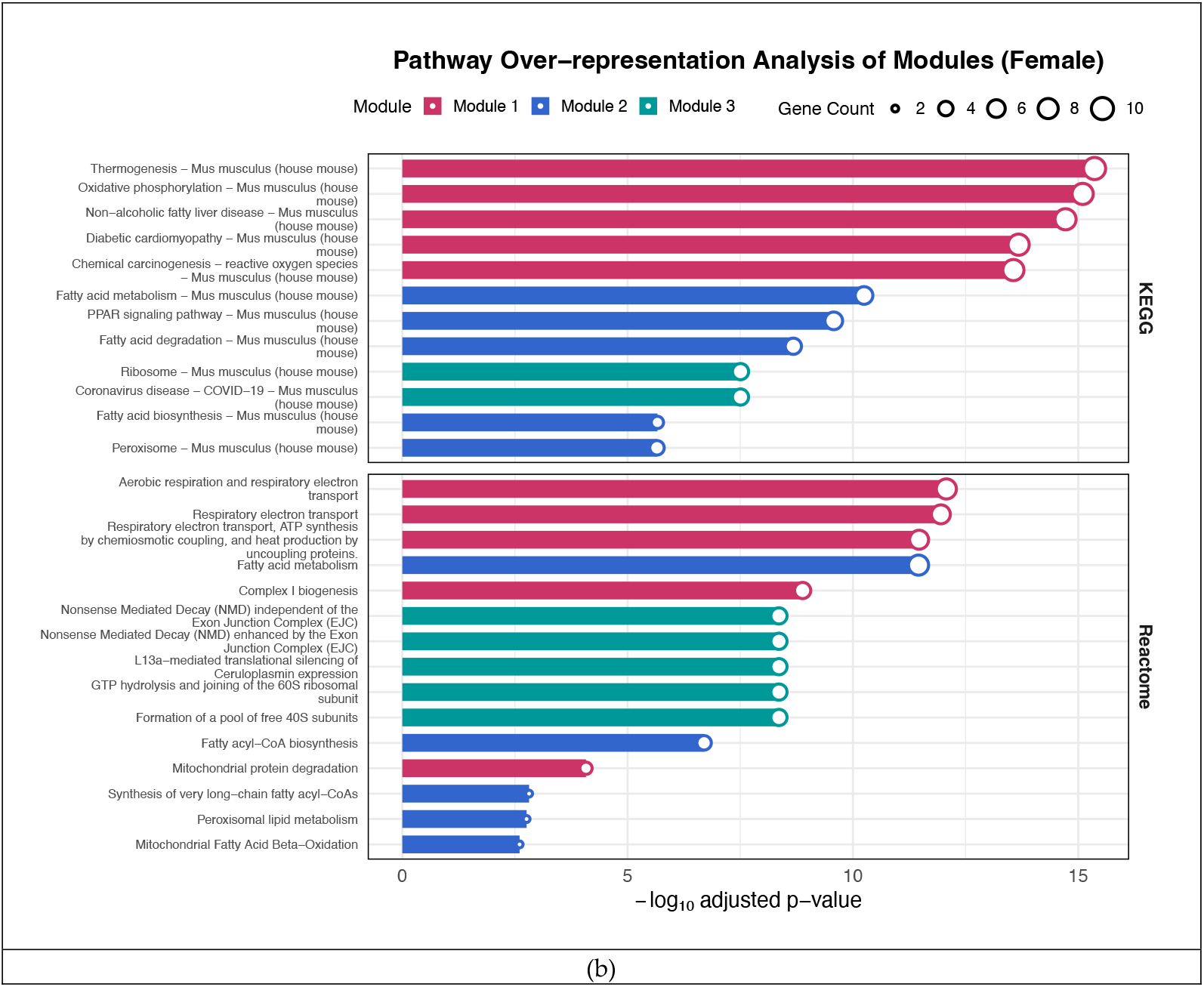
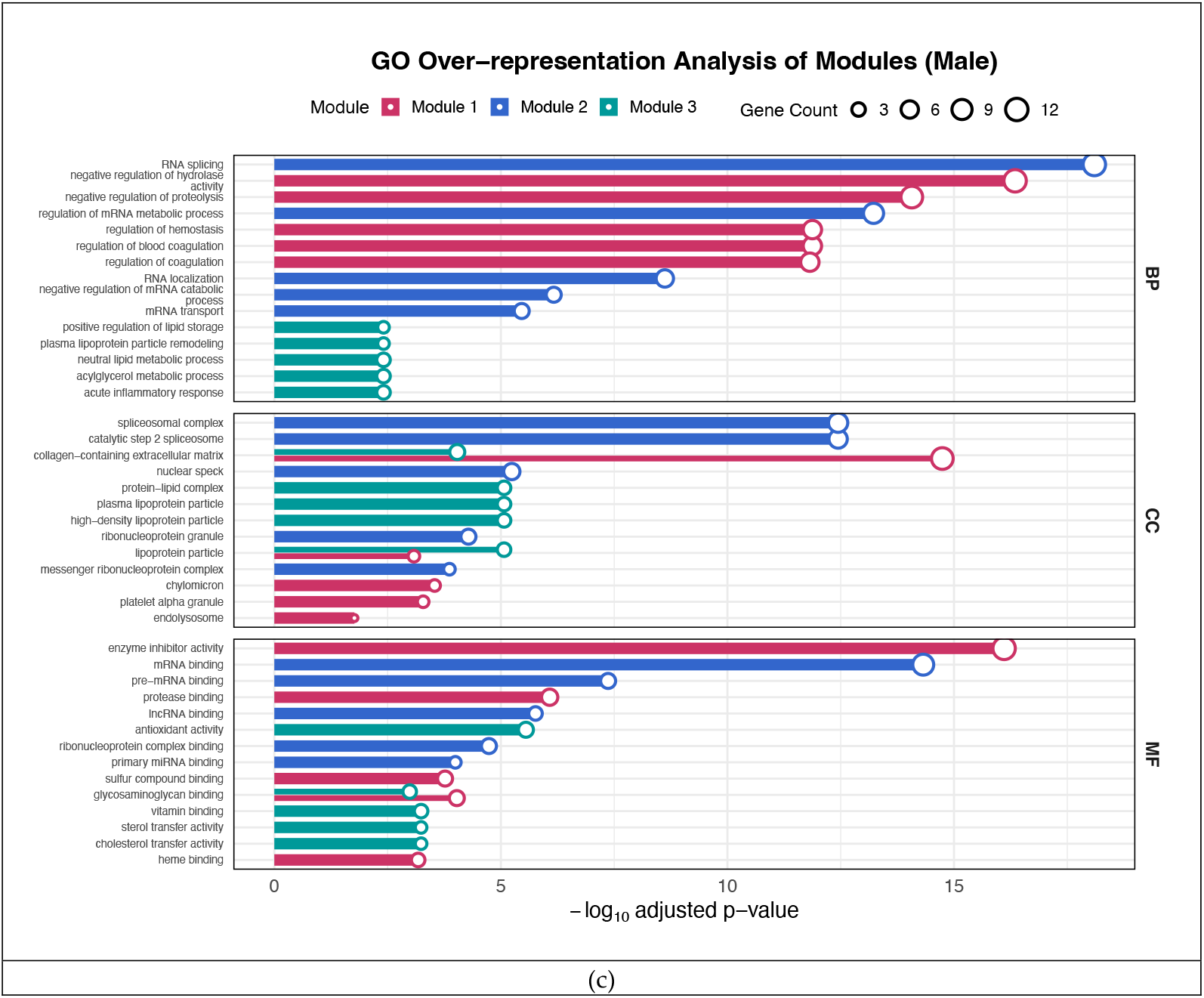
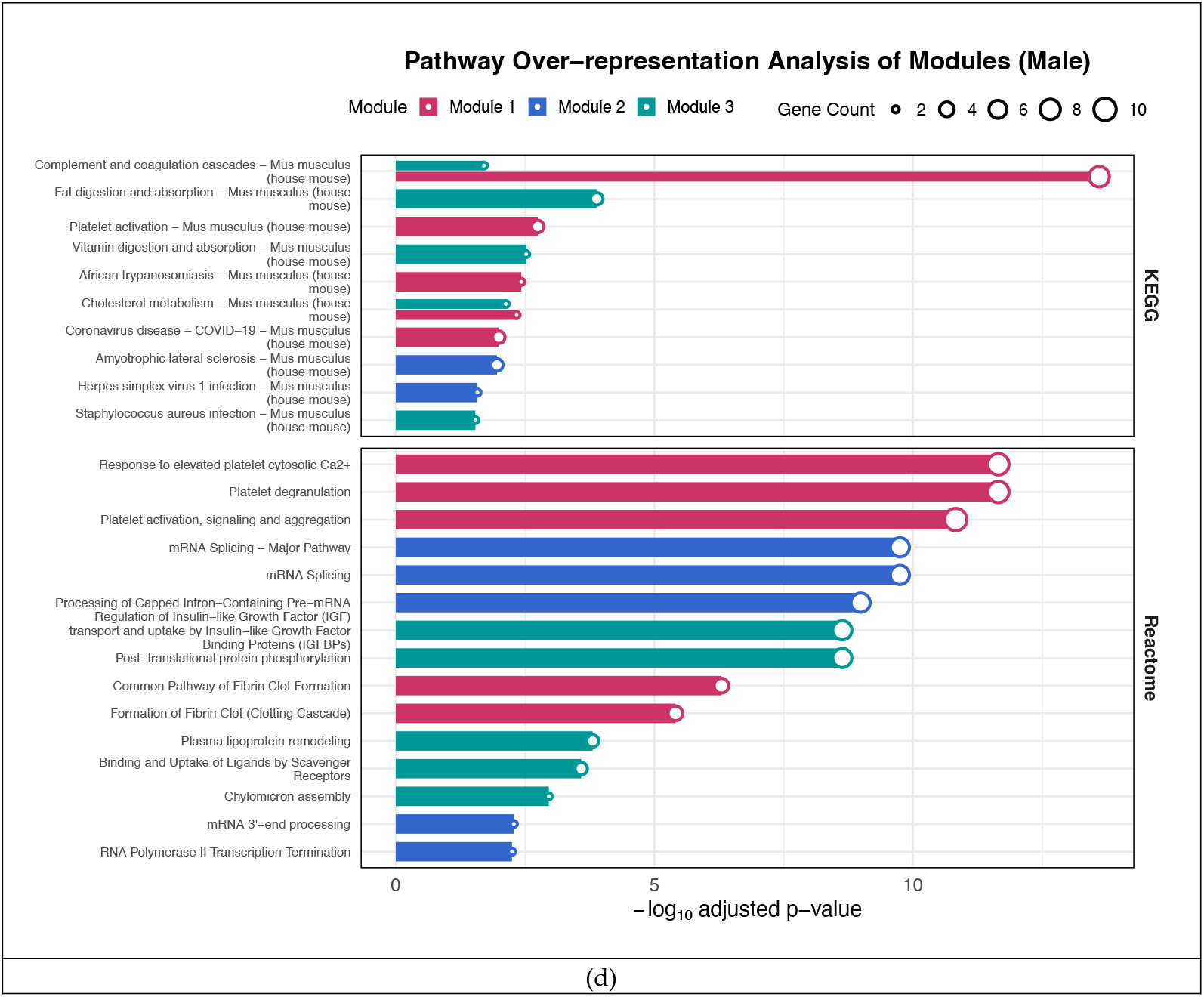
Overrepresentation analysis of the top three moules. Red bars represent module one, blue bars re-present module two and green bars represent module three:(**a, c)** The top significant terms of biological processes, cellular composition and molecular function from GO analysis of female and male groups *P*.adj < 0.05;(**b,d)** The top pathways from KEGG and reactome pathway analysis of top three modules of female and male mice.

GO term analysis of all DEPs showed enrichment in cortical cytoskeleton and actin filament binding processes (supplementary fig. 1).

Reactome pathway analysis further indicated the enrichment of mitochondrial respiratory pathways including respiratory electron transport, aerobic respiration and ATP synthesis by chemiosmotic coupling and other mitochondrial processes including heat production by uncoupling proteins, complex I biogenesis, mitochondrial protein degradation, mitochondrial fatty acid beta oxidation. Pathways associated with GTP hydroly-sis and joining of 60S ribosomal subunit and formation of a pool 40S subunit were also enriched (Figure 4b).

Oxidative phosphorylation and fatty acid metabolism were found to be enriched in both GO and KEGG analysis. KEGG pathway analysis also additionally represented enrichment of pathways of chemical carcinogenesis, fatty acid biosynthesis and degradation (Figure 4 a, b).

### 3.4. Cholesterol metabolism and Coagulation pathways in Male Mice

In GO term analysis of male mice, top clusters in the PPI network were found to be enriched in regulation of coagulation, positive regulation of lipid storage, plasma lipoprotein remodeling, neutral lipid metabolic process, acylglycerol metabolic process, RNA splicing, spliceosomal complex, catalytic step 2 spliceosome, regulation of mRNA metabolic process, mRNA transport, mRNA binding and RNA localization (Figure 4c). GO term analysis of all DEPs showed enrichment cortical cytoskeleton, actin filament and actin binding (Supplementary figure).

Reactome analysis also confirmed enrichment of platelet activation, signaling and aggregation, platelet degranulation, formation of fibrin clot and clotting cascade, mRNA processing pathways, Post™translational protein phosphorylation (Figure 4d).

KEGG pathway analysis confirmed enrichment in complement and coagulation cascade, fat digestion and absorption, platelet activation and cholesterol metabolism pathways (Figure 4d). Enrichment analysis of all DEPs showed enrichment in cytoskeleton, regulation of actin cytoskeleton pathways

### 3.5. Prognostic value of coagulation and cholesterol proteins in Male Mice and mitochondrial Protein NDUFA8 in Female mice

To identify the relationship between expression of Hub protein and overall survival time of patients and to evaluate prognostic value of Hub proteins, survival analysis of human homologs of all Hub genes was carried out using online tool UCLAN. The results indicated that eight hub genes were significantly associated with overall patient survival. Among the hub genes of DEN-treated female mice only the upregulation of NDUFA8 (P< 0.017) and ATP5H (P < 0.0066) was associated with poor patient survival (Figure 5 a, b). In DEN-treated male mice upregulation of FGG (P < 0.037), HPX (P < 0.016), FGA (P < 0.003), SERPINC1 (P < 0.023), APOA1 (P < 0.035) and HRG (P < 0.032) was associated with poor patient survival (Figure 5 c-h).

**Figure 5.**
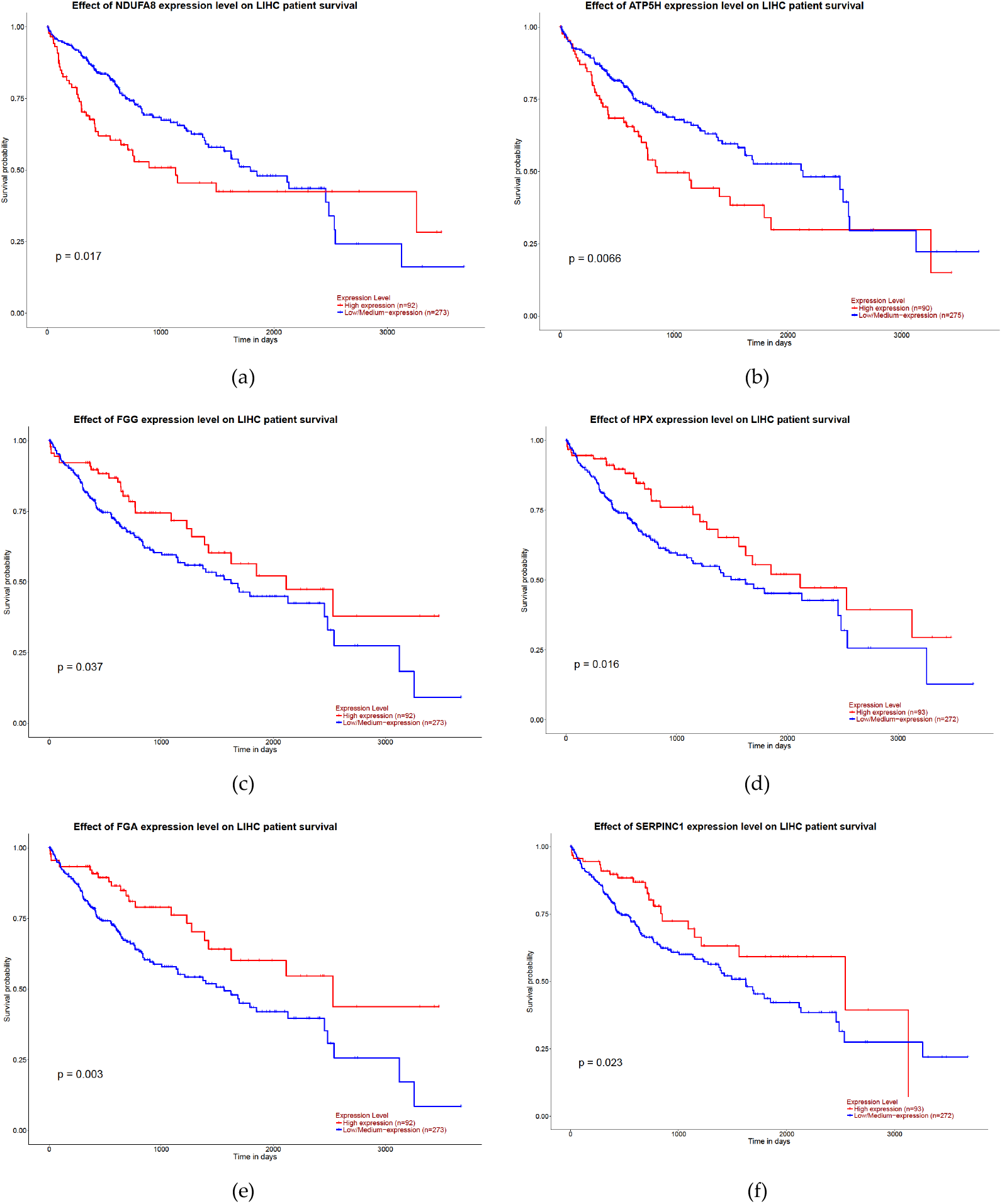
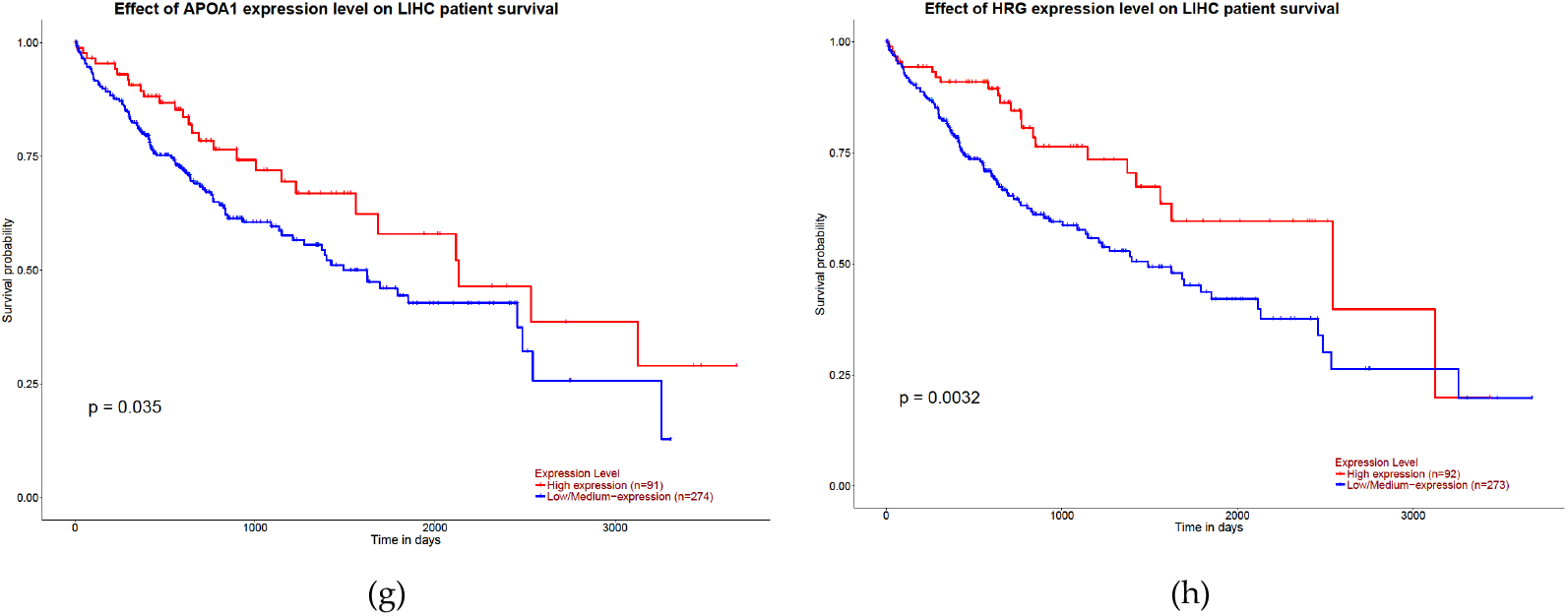
Overall survival curves for eight proteins with significant prognostic value: **(a)** NDUFA8**;(b)** ATP5H**; (c)** FGG**; (d)** HPX**; (e)** FGA**; (f)** SERPINC1**; (g)** APOA1**; (h)** HRG.

## 4. Discussion

In the present study, differential proteomic analysis of DEN-induced male and female HCC mouse models provided insights into gender specific proteomic signatures and associated molecular mechanisms. Two novel proteins (NDUFA8, ATP5H) in the female cohort and six proteins (FGG, FGA, HPX, SERPINC1, APOA1, HRG) in the male cohort were significantly associated with poor prognosis. The study provides comprehensive proteomic landscape of mouse HCC that emphasizes the need of using gender specific mouse models in HCC research.

While the study offers important insights, certain limitations should be acknowledged. Early mortality in male mice reduced statistical power for detecting subtle proteomic changes. To address this, we supplemented our dataset with proteomic data from eight DEN and CCl_4_-treated male mice from Zhang et al. [8]. Despite these constraints, our proteomic findings were internally consistent and validated through literature cross-referencing. However, due to resource limitations, we were unable to experimentally confirm novel proteins via techniques such as Western blotting or immunohistochemistry. Nevertheless, the use of integrative and literature-supported validation strengthens the reliability of our observations and lays the foundation for future targeted experimental studies.

Our findings build on and extend prior work by Sun et al. [15] and Zhang et al., [8] which investigated transcriptomic and proteomic changes in DEN and CCl_4_-induced HCC mouse models. Sun et al. [15] identified enrichment of pathways such as steroid hormone biosynthesis, retinol metabolism, and cytochrome P450 via transcriptomic profiling. Zhang et al. [8] reported 4,383 quantified proteins and 432 differentially expressed proteins (DEPs) in male mice, with upregulation of actin cytoskeleton components linked to tumor progression. While these studies provided foundational insights, Sun et al. [15] focused solely on transcriptomics, and Zhang et al. [8] limited their proteomic analysis to male mice. Our study complements and expands on these works by providing sex-specific proteomic data, capturing both overlapping and distinct biological pathways.

In the female cohort, we observed significant upregulation of oxidative phosphorylation and downregulation of fatty acid metabolism and xenobiotic detoxification. We found enrichment of mitochondrial respiratory chain components, particularly Complex I subunits such as NDUFA2, NDUFA7, NDUFA8, NDUFS6, NDUFV3, and *ATP5H*. Two of these, NDUFA8 and ATP5H were associated with poor survival in human HCC patients. NDUFA8, a critical Complex I subunit, regulates electron transport and ATP production and has been implicated in oncogenic proliferation in cervical cancers [16]. Complex I, comprising over 40 subunits, initiates the mitochondrial electron transport chain and is the largest and first point of electron entry for NADH oxidation [17]. While mitochondrial metabolism has been increasingly recognized in cancer biology, the role of Complex I subunits in hepatocellular carcinoma remains poorly characterized [18]. The observed mitochondrial shift toward enhanced respiratory metabolism may reflect an adaptive oncogenic response to sustain energy production and redox balance in tumor cells [18,19].

Concurrently, we observed the downregulation of fatty acid metabolism and xenobiotic detoxification pathways, particularly those mediated by CYP450 enzymes and chemical carcinogenesis–DNA adduct formation. This suppression may impair the liver’s intrinsic ability to metabolize and eliminate carcinogens, thereby allowing for ROS accumulation and prolonged genotoxic stress [20]. Elevated oxidative phosphorylation, combined with reduced detoxification capacity, may promote ROS buildup—further supported by our observation of upregulated ROS-associated signaling. This pro-oxidative environment can enhance DNA damage, impair lipid homeostasis, and promote tumorigenesis [21]. Downregulation of fatty acid β-oxidation, as reported in several HCC subtypes [22], likely contributes to lipid accumulation and metabolic stress, compounding the oncogenic effect. Collectively, our results suggest that female HCC may harbor a unique metabolic phenotype characterized by mitochondrial overactivation and impaired hepatic detoxification, which could represent a therapeutic vulnerability for sex-specific interventions targeting mitochondrial bioenergetics and ROS regulation.

In the male cohort of DEN-induced hepatocellular carcinoma (HCC), proteomic analysis revealed significant upregulation of pathways related to the regulation of the actin cytoskeleton, particularly in muscle cells, alongside enhanced cholesterol metabolism, as shown by KEGG pathway enrichment. Gene Ontology (GO) analysis further supported overrepresentation of processes related to lipoprotein particles, lipid storage, and lipid remodeling, indicating active metabolic reprogramming in tumor tissues. Several hub proteins identified in this group—including FGG, FGA, SERPINC1, HPX, ITIH4, PLG and HRG are associated with lipid transport, coagulation, and immune response regulation [23,24]. Many of these proteins also demonstrated prognostic significance: high expression of FGG, FGA, HPX, and SERPINC1 was associated with poorer overall survival [25,26].

The concurrent enrichment of coagulation-related proteins (e.g., FGG, FGA, SERPINC1) and cholesterol metabolism pathways points to a potential interaction between hemostatic and lipid regulatory systems in male HCC progression. Fibrinogen subunits, particularly FGG and FGA, are well known for their roles in blood clotting but have also been implicated in tumor invasion, metastasis, and angiogenesis via modulation of the epithelial–mesenchymal transition (EMT) [27–29]. Additionally, SERPINC1 (antithrombin III), beyond its anticoagulant role, has been reported to suppress tumor progression via modulation of the ubiquitin–proteasome system, contributing to apoptosis regulation and antitumor immunity [30]. These findings underscore a complex interplay between lipid regulation, coagulation, and immune function that contributes to male-specific HCC progression, offering novel therapeutic targets for personalized interventions.

In conclusion, this study presents a comprehensive proteomic characterization of male and female DEN-induced HCC mouse models, revealing distinct sex-specific mo-lecular signatures with implications for tumor progression and prognosis. While female HCC was marked by enhanced oxidative phosphorylation and suppressed detoxification pathways, male HCC exhibited upregulation of cholesterol metabolism and coagulation-associated proteins, suggesting divergent metabolic and immune microenvironments. Despite limitations such as early mortality in male mice and the translational gap between mouse and human models, the literature validation supports the biological relevance of our findings.

Future studies should include functional validation of hub proteins and their impact on sex-informed therapeutic strategies. This work lays a foundation for more personalized approaches in liver cancer research and may inform future clinical trial designs.

## Supporting information

GO analysis of female DEPs

GO analysis of male DEPs

KEGG and Reactome analysis of female DEPs

KEGG and Reactome analysis of male DEPs

## Supplementary Materials

The supporting information can be downloaded.

## Author Contributions

Conceptualization, Salmma Salamah Salihah Jamila Iqbal and Sana Mahmood; methodology, Salmma Salmah Salihah, Muhammad Tahir, Rabia Sultan and Bareera Bibi; software, Bareera Bibi and Rabia Sultan ; validation, Bareera Bibi; resources, Asma Gul, Munazza Raza Mirza, Muhammad Rizwan Alam and Martin R. Larsen; writing—original draft preparation, Salmma Salamah Salihah; writing—review and editing, Muhammad Tahir, Jamila Iqbal and Muhammad Rizwan Alam; visualization, Bareera Bibi; supervision, Jamila Iqbal, Asma Gul.; project administration, Asma Gul and Martin R. Larsen.; funding acquisition, Asma Gul. All authors have read and agreed to the published version of the manuscript.

## Institutional Review Board Statement

The animal study protocol was approved by the Institutional Review Board (or Ethics Committee) of International Islamic University, Islamabad (April 02, 2021) for studies involving mice.

## Informed Consent Statement

Not applicable.

## Data Availability Statement

Data is provided within the manuscript or supplementary files.

## Acknowledgments

We are grateful to the Pakistan Science Foundation for partially funding this research via PSF grant# 733. We are thankful to Molecular Oncology Research Lab for serving as a base camp for this project. We also appreciate Dr. Ikramullah for allowing us to use the animal house facility at the Center for Interdisciplinary Research in Basic Science (CIRBS), International Islamic University Islamabad. The study was supported by the VILLUM Center for Bio analytical Sciences at University of Southern Denmark and by PRO-MS, Danish National Mass Spectrometry Platform for Functional Proteomics.

## Disclaimer/Publisher’s Note

The statements, opinions and data contained in all publications are solely those of the individual author(s) and contributor(s) and not of MDPI and/or the editor(s). MDPI and/or the editor(s) disclaim responsibility for any injury to people or property resulting from any ideas, methods, instructions or products referred to in the content.

