## Supplementary figures and images for "Differential Proteomic Analysis of DEN-Induced Hepatocellular Carcinoma in Male and Female Balb/c Mice Reveals Novel Gender Specific Markers"

### GO analysis of female DEPs

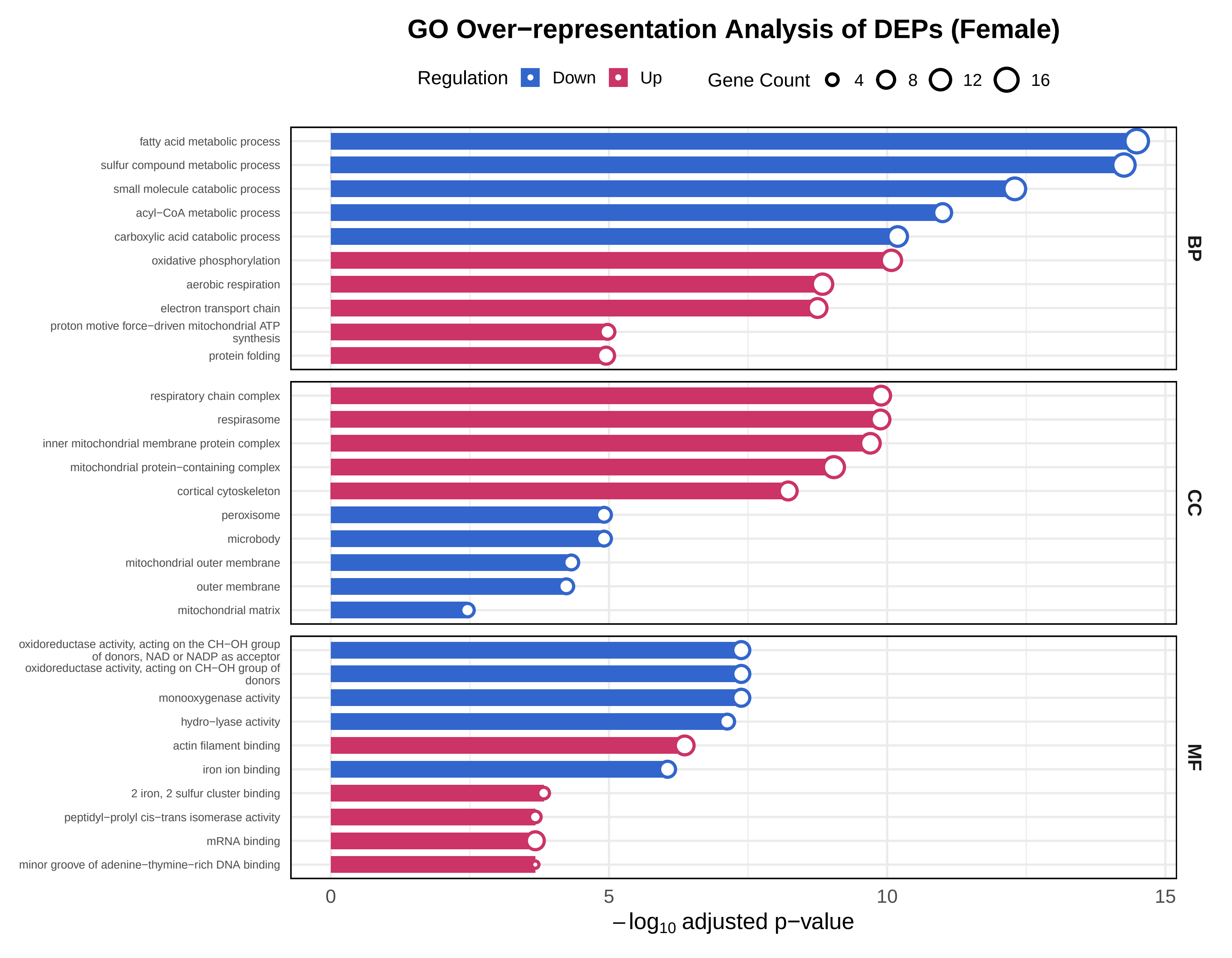

### GO analysis of male DEPs

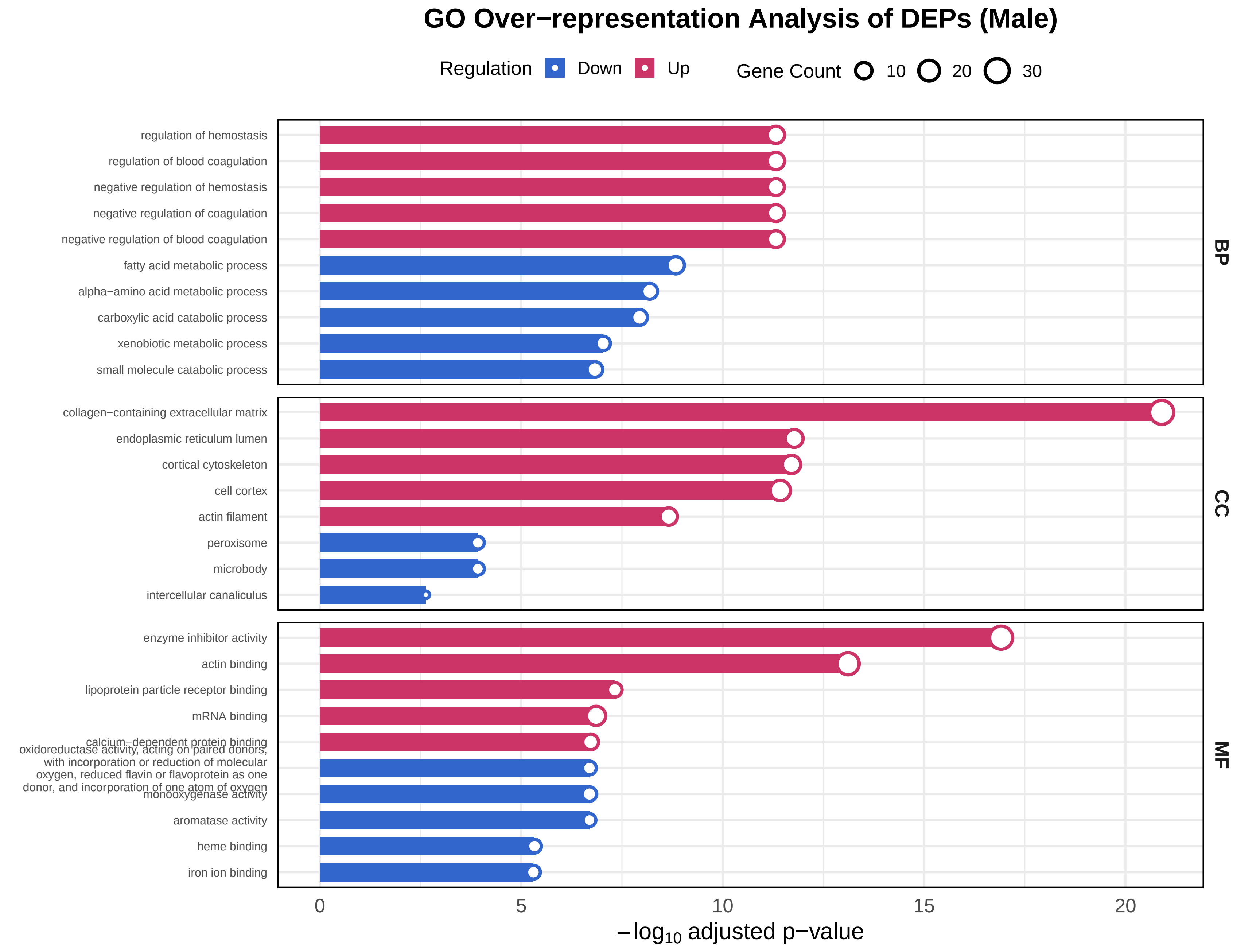

### KEGG and Reactome analysis of female DEPs

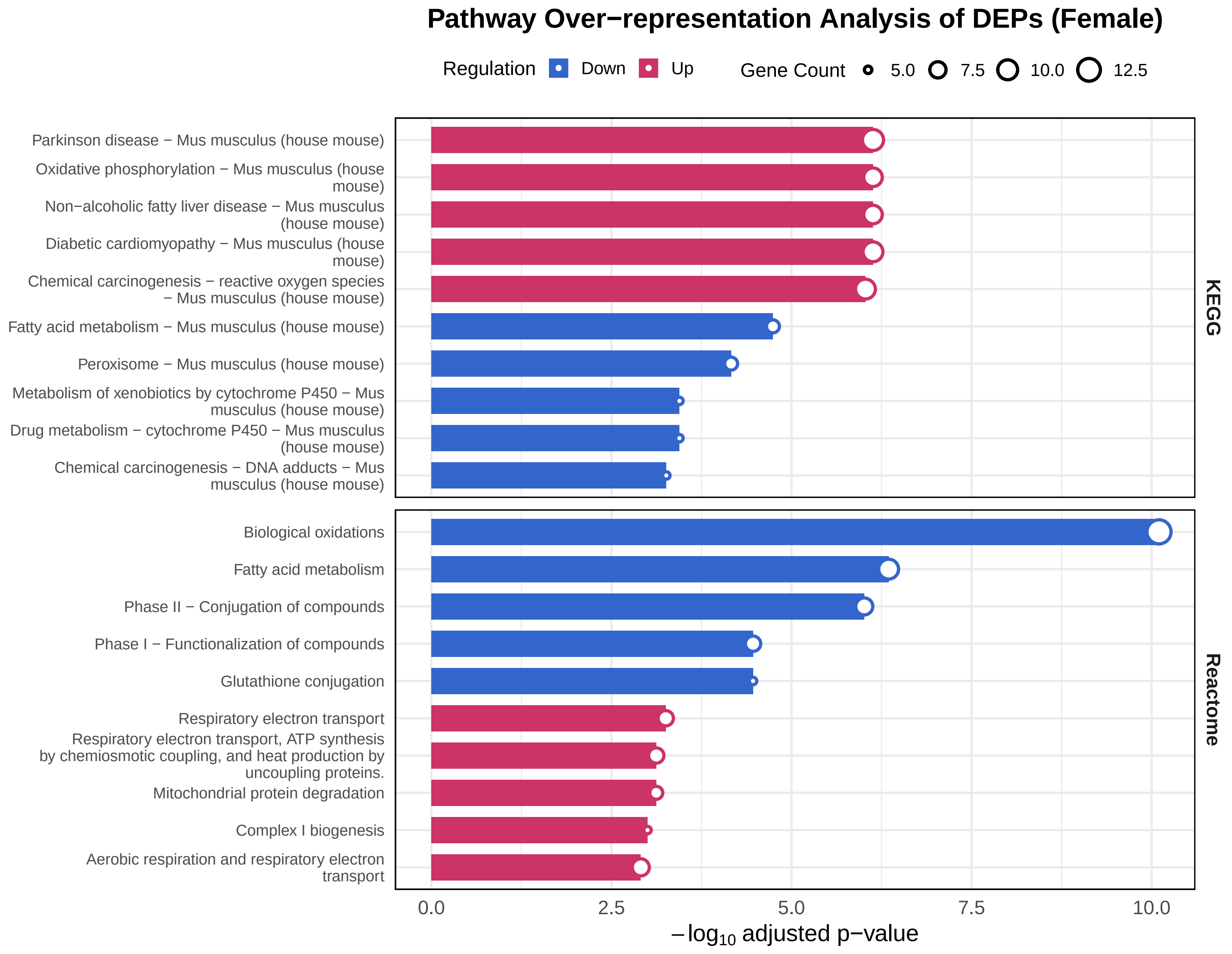

### KEGG and Reactome analysis of male DEPs

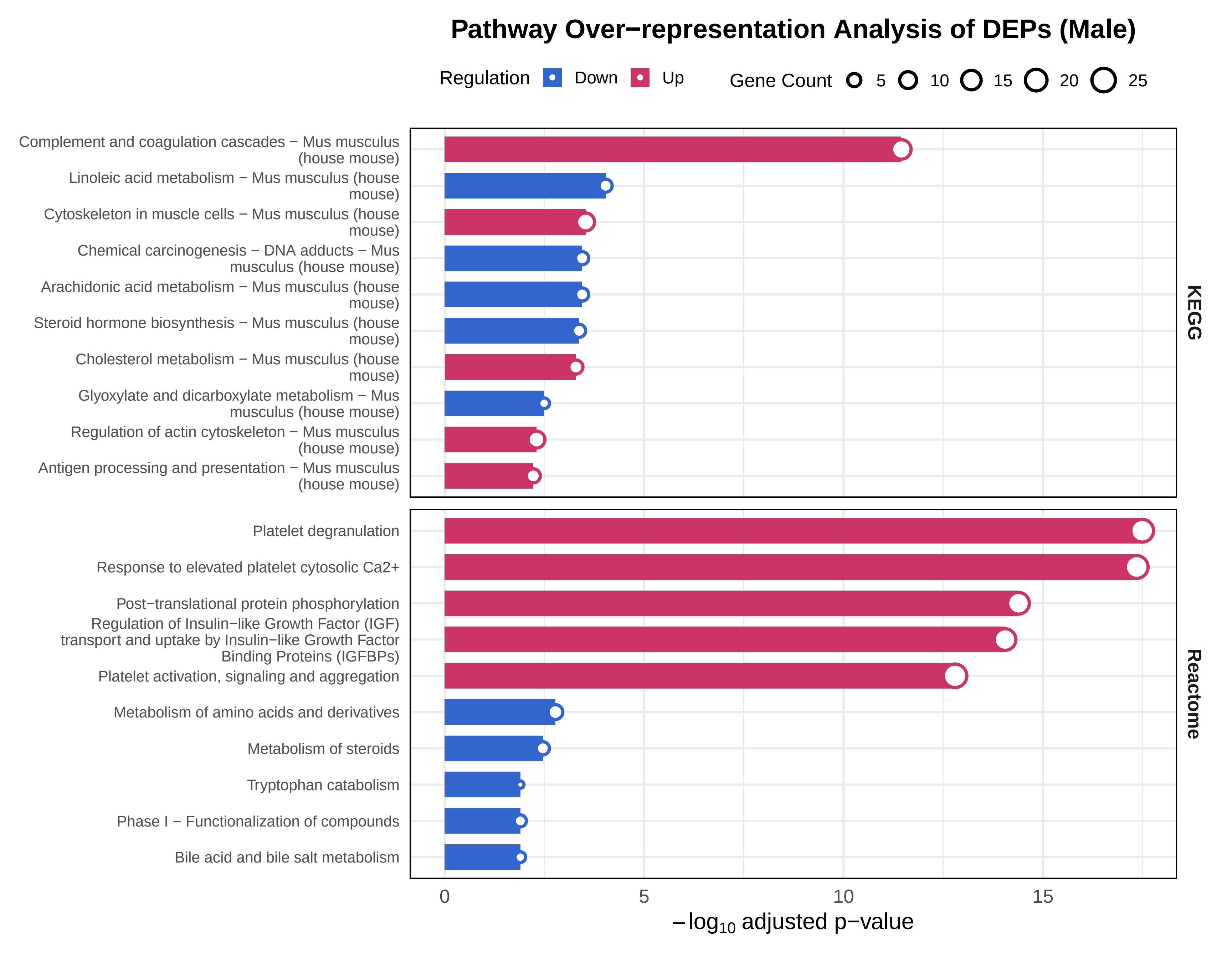
